# iDESC: Identifying differential expression in single-cell RNA sequencing data with multiple subjects

**DOI:** 10.1101/2022.02.07.479293

**Authors:** Yunqing Liu, Ningya Wang, Taylor S. Adams, Jonas C. Schupp, Weimiao Wu, John E. McDonough, Geoffrey L. Chupp, Naftali Kaminski, Zuoheng Wang, Xiting Yan

## Abstract

Single-cell RNA sequencing (scRNA-seq) enables assessment of transcriptome-wide changes at single-cell resolution. However, dominant subject effect in scRNA-seq datasets with multiple subjects severely confounds cell-type-specific differential expression (DE) analysis. We developed iDESC to separate subject effect from disease effect with consideration of dropouts to identify DE genes. iDESC was shown to have well-controlled type I error and high power compared to existing methods and obtained the best consistency between datasets and disease relevance in two scRNA-seq datasets from same disease, suggesting the importance of considering subject effect and dropouts in the DE analysis of scRNA-seq data with multiple subjects.

## Background

Recent advances in droplet-based single-cell RNA sequencing (scRNA-seq) technology have enabled investigators to assess transcriptome-wide differences at single-cell resolution [1-3]. Instead of pooling RNAs from all cells together, droplet-based scRNA-seq technology isolates cells using oil droplets, in which each cell is lysed and a cell barcode and a unique molecular identifier (UMI) are added onto the amplified cDNAs. Using these cell barcodes and UMIs, sequencing reads are demultiplexed into different cells and transcripts, which enables single-cell transcriptome profiling without PCR amplification bias. In recent years, scRNA-seq has been used to study cellular heterogeneity and gene expression variability across different cell types in diverse human tissues [4] and diseases (chronic diseases [5], infectious diseases [6], autoimmune diseases [7], and cancers [8]). These applications have revealed disease-related cell type specific transcriptomic changes [9], rare cell types [10], and cell type composition changes [11], providing important insights into disease pathogenesis and facilitating the development of personalized treatment of diseases [12, 13].

Despite the great potential of scRNA-seq technology, challenges remain in the corresponding data analysis. Specifically, one common task in scRNA-seq data analysis is to identify cell type specific differentially expressed (DE) genes between two groups of subjects [14], which can be challenging due to prevalent dropout events and substantial subject effect, or so-called between-replicate variation [15]. Dropout refers to the event when a given gene is observed at a moderate expression level in one cell but is not detected in another cell of the same type from the same sample [16], leading to underestimation of gene expression level and overestimation of variation in the data which may generate false positive results. Moreover, with the popularity of multi-sample scRNA-seq datasets from different diseases, tissues, and cell types, many of them have consistently shown that within the same cell type, cells of the same subject cluster together but separate well from cells of other subjects [6, 17, 18]. This suggests that there exists a large variation across subjects possibly due to heterogeneous genetic backgrounds or environmental exposures and this variation is much larger than the within-subject variation across cells. Dominant subject effect severely confounds the DE analysis in scRNA-seq data because genes driving differences across subjects are likely to also be significantly different between two groups [15, 19, 20]. Taken together, in the DE analysis of scRNA-seq data with multiple subjects, it is critical to separate subject effect from disease effect with consideration of dropout events.

Technical batch effect is one possible reason for the large variation across subjects because many studies processed cells and cDNA libraries from different subjects in different batches due to the requirement of sample freshness in certain tissue types and early-stage scRNA-seq protocols. This may lead to batch effect in the data so that cells of the same type from different subjects have different expression profiles. However, in the scRNA-seq dataset from patients with idiopathic pulmonary fibrosis (IPF), the large variation across subjects was still present after adjusting for batch effect using scVI [21]. In addition, recent advances in combining scRNA-seq with upstream cell cryopreservation using dimethyl sulfoxide (DMSO) have enabled preservation of cells so that samples from different subjects can be processed together [22]. Comparison between scRNA-seq data from the same sputum sample with and without DMSO preservation showed no significant difference between the fresh and DMSO data but significant separation between different subjects was still present (unpublished data). Since the fresh and DMSO data from the same sample were generated in two different batches, this confirmed that the large between-subject variation was a consequence of biological subject effect but not technical batch effect. Therefore, it is inadequate to consider this variation as technical batch effect and remove it using batch effect adjustment tools in scRNA-seq data. In fact, removing this variation as technical batch effect may remove the disease effect of interest because subject effect confounds with disease effect.

Many DE analysis methods for scRNA-seq data have been developed and compared [23-25]. They can be classified into two categories depending on whether subject effect is considered. Although methods that ignore subject effect have been used in DE analysis, they are more suitable for identification of marker genes for a given cell type, which is fundamentally different from DE analysis.

Within the category of methods that ignore subject effect, there are methods specifically designed for scRNA-seq data and methods adopted from bulk RNA-seq DE analysis. Among the methods designed for scRNA-seq data, BASiCS [26] and TASC [27] require external RNA spike-ins to provide information on technical variation and use a Bayesian hierarchical Poisson-Gamma model and a hierarchical Poisson-lognormal model, respectively, to fit data. Monocle [28-30] and NBID [31] model UMI counts of each gene using a negative binomial distribution without considering dropouts. To account for dropouts, a group of methods were developed including DEsingle [32], DESCENT [33], SC2P [34], SCDE [16] and MAST [35]. These methods utilize mixture models or hierarchical models, mostly zero-inflated, to model dropouts and captured transcripts. DEsingle fits a zero-inflated negative binomial model in each group and conducts a likelihood ratio test for significance assessment. DESCENT models UMI counts using a hierarchical model assuming that the true underlying expression follows a zero-inflated negative binomial distribution and the capturing process generating the observed data follows a beta-binomial distribution. SC2P models dropout events using a zero-inflated Poisson distribution and fits the detected transcripts using a lognormal-Poisson distribution. The assumption in SC2P that the cell-specific dropout rate and dropout distribution are shared by all genes may eliminate the natural stochasticity in scRNA-seq data. SCDE employs a two-component mixture model with a negative binomial and a low-magnitude Poisson component to model efficiently amplified read-outs and dropout events, respectively. The dropout rate for a given gene is determined by its true underlying expression level in the cell, which is estimated based on a selected subset of highly expressed genes. MAST uses a two-part hurdle model in which dropout rates are modeled by a logistic regression model and non-zero expression follows a Gaussian distribution. SC2P, SCDE and MAST were originally designed for Transcript Per Kilobase Million (TPM) data which has different technical noise and data distribution from UMI count data [36]. Multimodality has been observed in scRNA-seq data due to cellular heterogeneity within the same cell type. To consider multimodality, scDD [37] was designed to model count data with a Dirichlet process to detect genes with difference in mean expression, proportion of the same component, or modality between groups. D3E [38], a nonparametric method, fits a bursting model for transcriptional regulation and compares the gene expression distribution between two groups. It was previously reported to generate false-positive results on negative control datasets [24]. A recent study [23] showed that bulk RNA-seq analysis methods, including DESeq2 [39], limma-trend [40], and Wilcoxon rank sum test [41], have comparable performance to methods designed for scRNA-seq data when applied to the cell-level UMI count data, especially after filtering out lowly expressed genes.

All the methods mentioned above treat cells from the same subject as independent, which may be efficient for identifying cell type marker genes, but inappropriate for DE analysis to identify disease or phenotype associated genes due to the presence of dominant subject effect confounded with disease effect as described above. One simple and straightforward solution is to aggregate expression levels of cells from the same subject by averaging and then to compare the aggregated sample-level “pseudo-bulk” expression levels between two groups of subjects using Student’s t test. We denote this method as subject-t-test (subT). Furthermore, two recent studies proposed the following three DE analysis methods to consider subject effect in scRNA-seq data. Zimmerman et al. [19] developed MAST-RE by adding a subject random effect to the non-zero expression part of the hurdle model in MAST. The muscat package [20] provides two approaches to consider subject effect: (1) muscat-PB that aggregates cell-level UMI counts into sample-level “pseudo-bulk” counts which are then compared between two groups using edgeR that was developed for DE analysis in bulk RNA-seq data; and (2) muscat-MM that fits a generalized linear mixed model (GLMM) on the cell-level UMI counts to account for subject variation. Both muscat-PB and muscat-MM were compared to other methods and shown to have power gain by considering subject effect [15].

In this article, we develop a new statistical model to identify DE genes in scRNA-seq data with multiple subjects, named iDESC. A zero-inflated negative binomial mixed model is used to consider both subject effect and dropouts. iDESC models dropout events as inflated zeros and non-dropout events using a negative binomial distribution. In the negative binomial component, a random effect is used to separate subject effect from group effect. Wald statistic is used to assess the significance of group effect. We compared iDESC with some of the existing DE analysis methods based on type I error and statistical power in both simulated and real datasets.

## Results

### Method performance evaluation overview

We evaluated and compared the performance of iDESC and eleven existing DE analysis methods (Table 1) using both simulation studies and two real datasets. We divide these methods into two categories, based on whether subject effect is considered. The first category of methods considers subject effect, including iDESC, MAST-RE, muscat-PB, muscat-MM and subT. Among them, iDESC, MAST-RE and muscat-MM are mixed model-based methods, whereas muscat-PB and subT are aggregation-based methods. iDESC and MAST-RE also consider dropouts in scRNA-seq data. The other category of methods does not consider subject effect. Within this category, DEsingle, MAST and scDD consider dropouts while NBID, DESeq2, limma-trend and Wilcoxon rank sum test do not. DEsingle considers dropout in the fitted model but tests for differences in both group means and dropout rates instead of difference in group means only. Therefore, unlike most DE methods, DEsingle identifies genes with composite differences in group means and dropout rates. Comparison of these methods enabled us to assess the benefit of considering dropout evens and/or subject effect in the DE analysis of scRNA-seq data.

**Table 1.** Overview of the twelve DE analysis methods for comparison

|  | Dropout | Subject effect | Test | Model |
| --- | --- | --- | --- | --- |
| **iDESC** | √ | Mixed model | Wald test | Zero-inflated negative binomial mixed model |
| **MAST-RE** | √ | Mixed model | Likelihood ratio test | Two-part hurdle mixed model |
| **Muscat-MM** | × | Mixed model | Wald test | Negative binomial mixed model |
| **Muscat-PB** | × | Aggregation | Quasi-likelihood F-test | EdgeR on sample-level aggregated data |
| **subT** | × | Aggregation | Student's T test | T test on sample-level aggregated data |
| **DEsingle** | √ | × | Likelihood ratio test | Group-specific zero-inflated negative binomial model |
| **MAST** | √ | × | Likelihood ratio test | Two-part hurdle model |
| **scDD** | √ | × | Kolmogorov-Smirnov test | Dirichlet process mixture of normals |
| **NBID** | × | × | Likelihood ratio test | Negative binomial model with group-specific dispersion |
| **DESeq2** | × | × | Wald test | Negative binomial model with the same dispersion between groups |
| **limma** | × | × | Moderated T test | Linear regression model |
| **Wilcoxon** | × | × | Wilcoxon rank sum test | Nonparametric test |

We compared the method performance based on their type I error, statistical power, and consistency of the identified DE genes in two independent scRNA-seq datasets on the same disease, the Kaminski dataset [18] and the Kropski dataset [42]. Both datasets measured scRNA-seq data of whole lung tissue from patients with IPF and normal controls.

### Type I error assessment

To assess type I error, we permuted the group labels of subjects in both Kaminski and Kropski datasets 500 times. All twelve methods were applied to the permuted datasets to identify DE genes. For each gene, we calculated the empirical type I error as the proportion of permuted datasets with a p-value < 0.05 and compared it to the nominal level 0.05. Fig. 1 shows the empirical type I error of each method. In both datasets, methods that account for subject effect, including iDESC, MAST-RE, muscat-MM, muscat-PB and subT, had well-controlled type I error. Among these methods, MAST-RE had slightly inflated type I error for some genes, likely due to the deviation of log-normalized UMI counts from the assumed Gaussian distribution. In contrast, the type I error of the methods that do not consider subject effect were severely inflated. The inflation of type I error was more prominent in the Kropski dataset for these methods, indicating a larger subject effect in the data. Among these methods, DEsingle, MAST and scDD had the largest inflation in type I error. DESeq2 had the largest variation in type I error across all genes. Taken together, these results suggested that it is important to consider subject effect for type I error control in the DE analysis of scRNA-seq data with multiple subjects.

**Figure 1.**
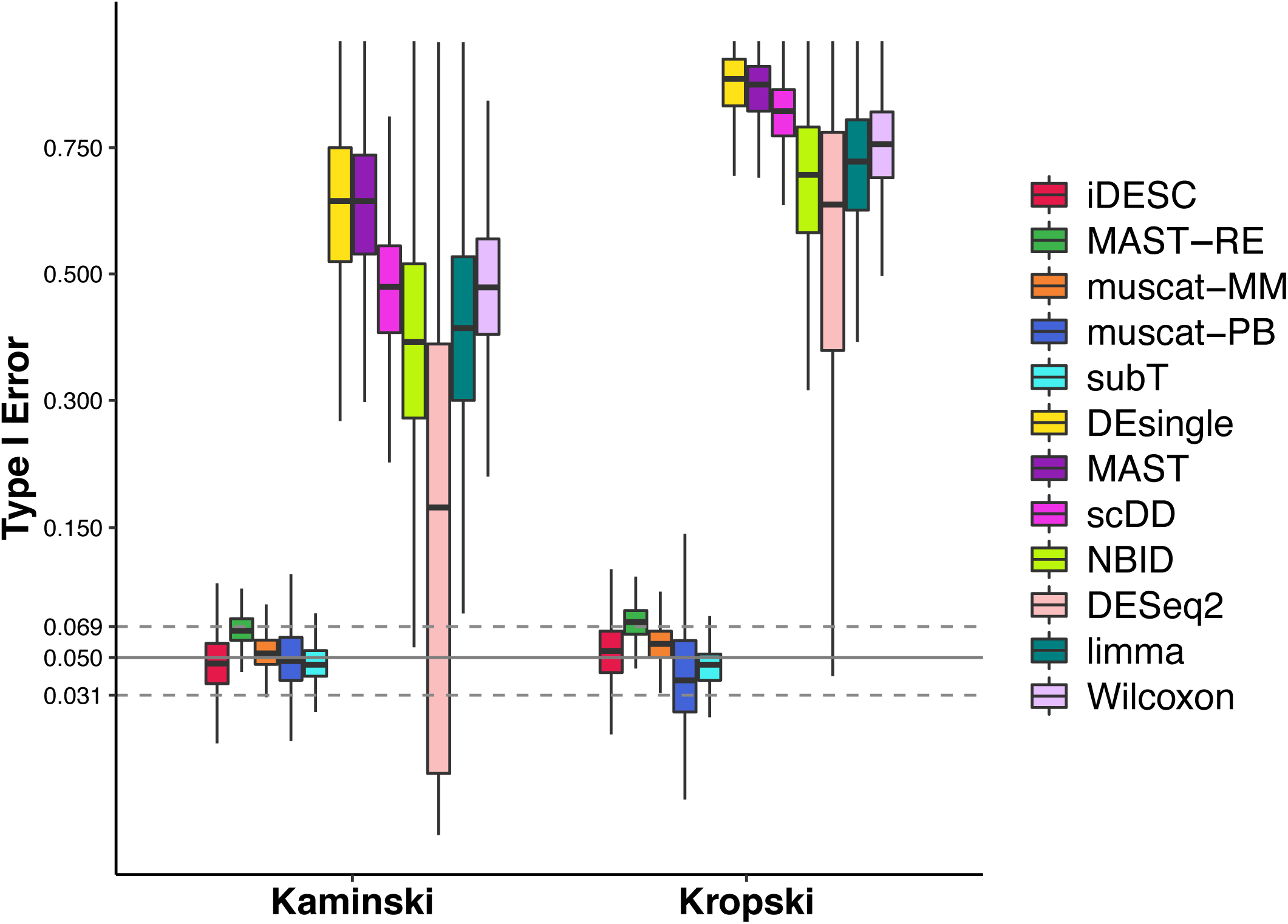
Empirical type I error of all methods on the two permuted real datasets. Boxplots show the median (center line), interquartile range (hinges), and 1.5 times the interquartile (whiskers) of empirical type I error at the nominal level of 0.05. Confidence interval of type I error is marked by two dashed lines (0.031-0.069).

### Power comparison

To compare power, we simulated scRNA-seq data under a wide range of parameter settings estimated from real datasets and conducted DE analysis using the twelve methods on the simulated data. Method performance was assessed by the area under a receiver operating characteristic curve (AUC) that describes the sensitivity and specificity of the identified DE genes under different significance levels where genes with non-zero group effect in the simulation model were treated as ground truth.

We considered two scenarios for dropout rates. In the first scenario where there was no dropout or dropout rates were the same between two groups, iDESC performed the best with the highest AUC in most of the simulation settings when compared to methods considering subject effect (Fig. 2a). subT had the second highest AUC but was slightly better than iDESC when the intercept was low (*α* = −9.3) and the group effect was positive and low (*β* = 0.1), corresponding to the situation of overall low expression in both groups and slightly higher expression in the disease group. The other three methods that account for subject effect, MAST-RE, muscat-MM and muscat-PB, had lower AUC than iDESC across all simulation settings. When we compared iDESC with the other seven methods that do not consider subject effect, Fig. S1a shows that iDESC achieved the highest AUC in detecting DE genes when the group effect was negative, i.e., the mean gene expression of the disease group was lower than that of the control group. However, when the group effect was positive, i.e., the mean gene expression of the disease group was higher than that of the control group, DEsingle, MAST, DESeq2, limma and Wilcoxon achieved improved power and had comparable or even higher AUC than iDESC. scDD and NBID had compromised performance across all simulation settings.

**Figure 2.**
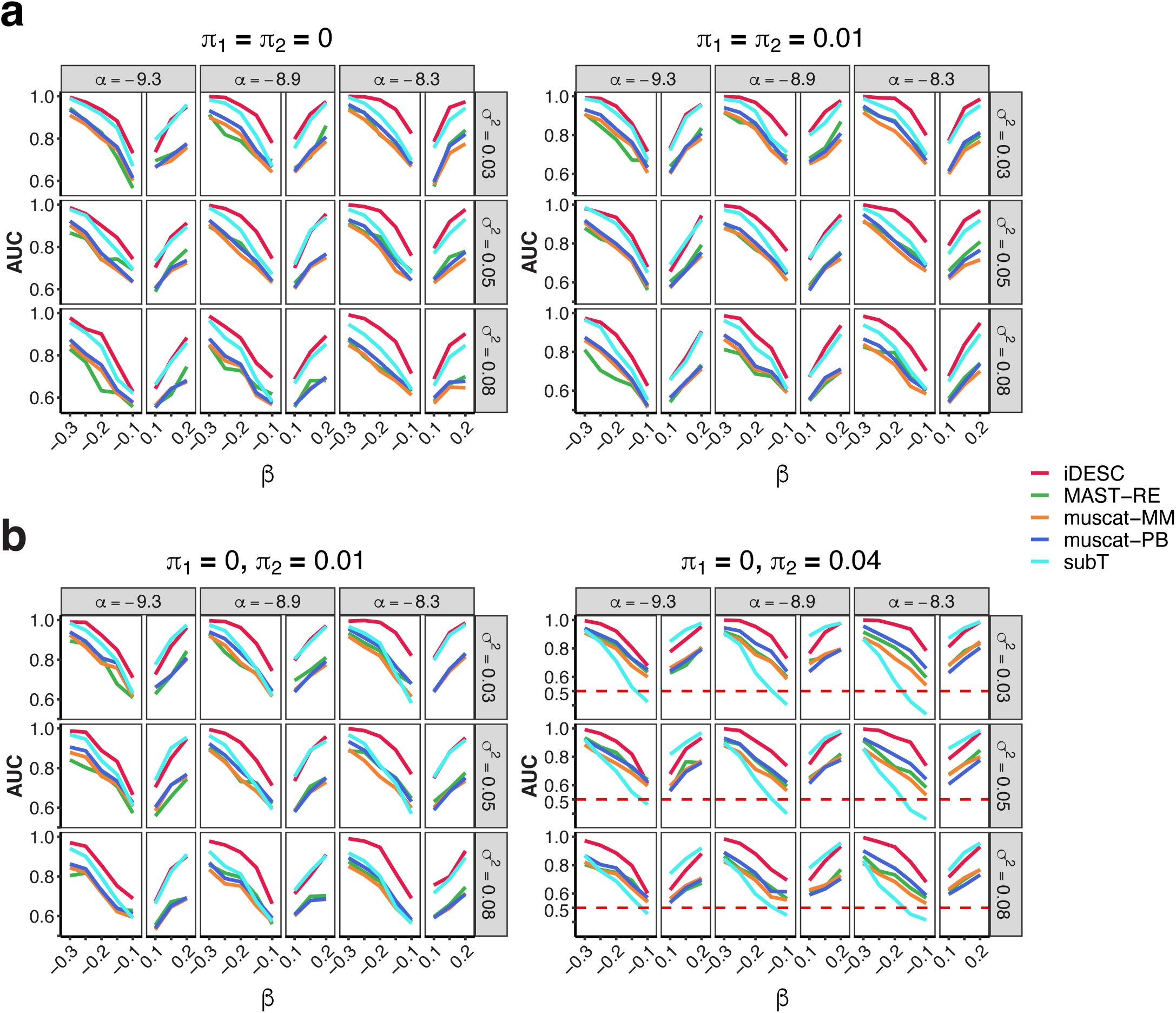
DE analysis accuracy of all methods in simulated datasets. Area under an ROC curve (AUC) was calculated to measure the accuracy of identified DE genes under two scenarios of dropout rate settings in disease (*π*_1_) and control (*π*_2_) groups: (A) Scenario I with the same dropout rate (0 and 0.01) between groups; and (B) Scenario II with dropout rate higher in the control group.

In the second scenario where the dropout rate was higher in the control group (*π*_2_ > *π*_1_), Fig. 2b shows that iDESC always had the highest AUC when the group effect was negative. subT had the largest decrease in performance among methods that account for subject effect when the difference in dropout rate between the two groups increases (*π*_1_ = 0, *π*_2_ = 0.04). However, when the group effect was positive, subT had improved performance and achieved comparable or even higher AUC than iDESC. This was expected because subT is designed to detect the observed mean expression difference between groups. When the mean gene expression of the disease group is higher and the dropout rate of the disease group is lower than those of the control group, the observed mean expression difference will be larger than the true difference between the two group means, facilitating subT to detect the expression difference between the two groups, especially when the difference in dropout rate is large and in an opposite direction to the group mean difference. In contrast, when the mean gene expression and the dropout rate of the disease group are both lower than those of the control group, the observed mean expression difference becomes smaller than the true difference so that subT loses power to detect the expression difference between the two groups, especially when the difference in dropout rate is large and in the same direction as the group mean difference. This suggests that subT is highly sensitive to difference in dropout rate between the two groups. The other three methods that account for subject effect, MAST-RE, muscat-MM and muscat-PB, had lower AUC than iDESC across all simulation settings. When we compared iDESC with the other seven methods that do not consider subject effect, Fig. S1b shows that iDESC achieved the highest AUC when the group effect was negative, while scDD had compromised power. However, when the group effect was positive, DEsingle, MAST, DESeq2, limma and Wilcoxon achieved improved power and had comparable or even higher AUC than iDESC, due to the same reason for similar behavior of subT in Fig. 2b described above. scDD and NBID had the worst performance. The overall poor performance of scDD in both scenarios may be attributed to the fact that scDD is designed for data with multimodality to detect difference in mean expression, proportion of the same component, or modality between groups.

In summary, iDESC had the highest AUC except for the settings with positive group effect and higher dropout rate in the control group where the observed difference is larger than the true group mean difference which favor the methods that ignore dropouts.

### Consistency of results in two scRNA-seq datasets

We used two independent scRNA-seq datasets of whole lung tissue from IPF patients and normal controls [18, 42], the Kaminski and Krospki datasets, to demonstrate and compare the performance of different methods. Since macrophage-driven transcriptomic changes in IPF patients have been previously reported [43-48], we focused on macrophages in both datasets and applied all five methods that consider subject effect, iDESC, MAST-RE, muscat-MM, muscat-PB and subT. After data processing and gene filtering, we had 6,564 genes and 21,433 macrophages in the Kaminski dataset, and 6,128 genes and 10,520 macrophages in the Kropski dataset. Macrophages from these two datasets were integrated using Seurat (Fig. 3a) and selected so that they are transcriptionally and functionally similar macrophages, ensuring their corresponding DE analysis results to be biologically comparable. At the threshold of false discovery rate (FDR) < 0.1, iDESC identified 2,462 and 239 DE genes in the Kaminski and Kropski datasets, respectively. The top upregulated DE genes in IPF such as *CTSK, FN*1, *SPP*1, *CCL*18 (Fig. 3b), were previously reported to be upregulated in IPF and related to IPF pathogenesis in macrophages [17, 18, 44, 45, 47-49]. To examine subject variation in these two datasets, we estimated cell-level effect coefficients [20] for each subject (Fig. 3c). Cell-level effect coefficients summarized the extent to which each cell reflects the group-level fold-change. Both inter- and intra-subject variations of effect coefficients in the Kropski dataset were larger than those in the Kaminski dataset, suggesting that the Kropski dataset had larger subject variation.

**Figure 3.**
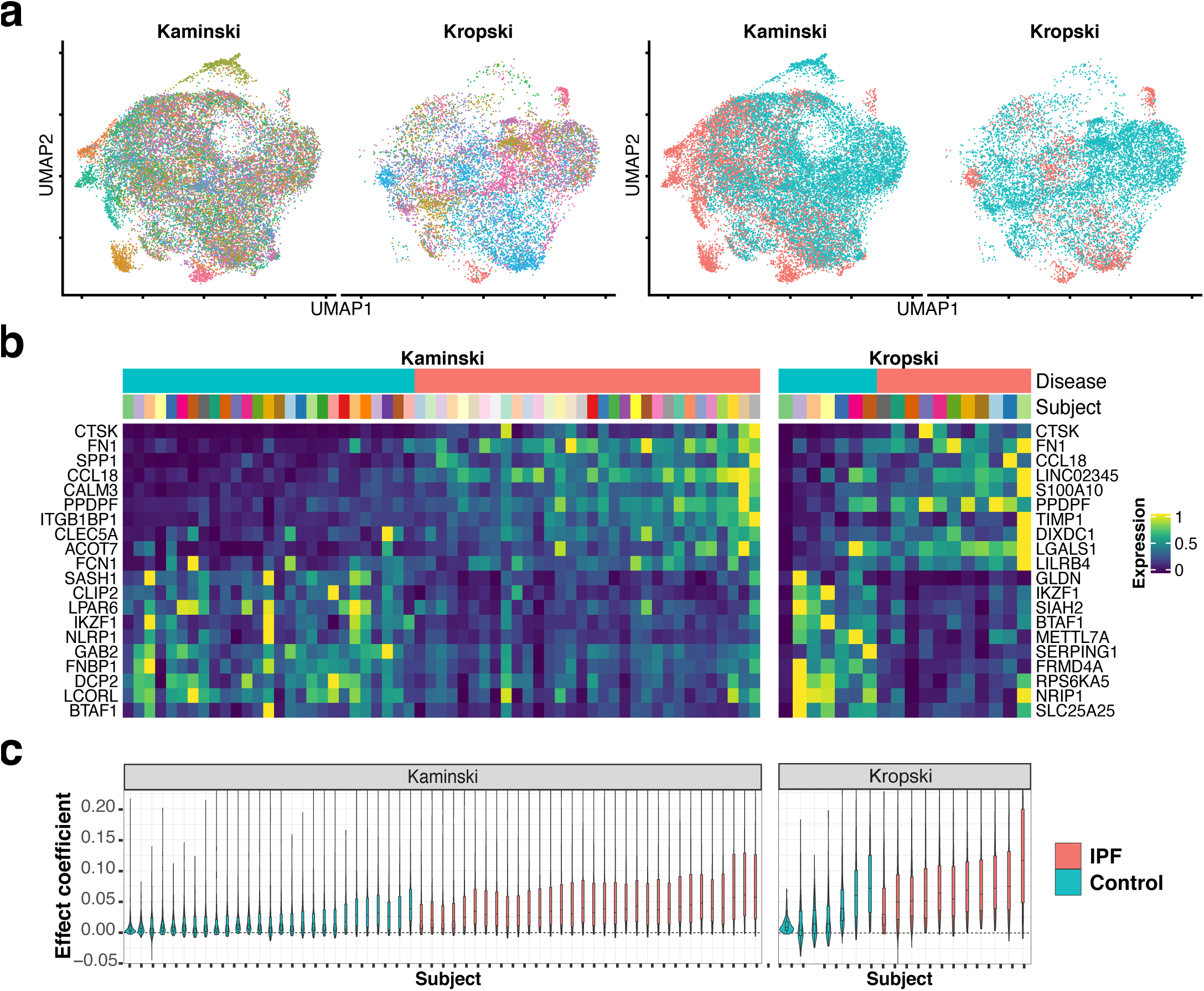
DE analysis using iDESC on two IPF macrophage datasets. A) UMAP of selected macrophages in the Kaminski and Kropski datasets colored by subject (left) and group (right). B) Heatmap of subject-level average expression for the top 10 upregulated and top 10 downregulated genes. C) Violin plots demonstrate cell-level contributions to the group fold-change within each subject. Each violin corresponds to one subject and is colored by group.

Fig. 4a shows the number of DE genes identified in the two datasets by each of the five methods. Due to larger subject variation in the Kropski dataset, all methods identified much less DE genes in the Kropski dataset than in the Kaminski dataset. iDESC identified the largest number of overlapping DE genes (123 genes) between the two datasets. We then evaluated method performance based on the consistency of DE genes between the two datasets. For each method, we conducted Fisher’s exact test to assess the significance of the overlap and calculated Jaccard index to measure the similarity between the two DE gene lists. The higher the overlap is, the more consistent and validated the results are between the two datasets, indicating a better performance. Table 2 shows that iDESC had the most significant overlap (p = 4.1×10^−6^), followed by muscat-PB (p = 7.2×10^−5^). In contrast, MAST-RE, muscat-MM and subT did not achieve significant overlap between the two datasets. In addition, iDESC had the largest Jaccard Index (JI = 0.076), followed by muscat-PB (JI = 0.016), while MAST-RE, muscat-MM and subT had much smaller Jaccard Index. Furthermore, we used a list of 83 IPF-related genes in the Harmonizome database [50] to validate the overlapping DE genes between the two datasets identified by each method. iDESC had the highest proportion of validation (7.23%), followed by muscat-PB (2.41%). None of the DE genes identified by muscat-MM and subT were validated. Taken together, iDESC performed the best and achieved the most consistent and validated results in real datasets.

**Table 2.**
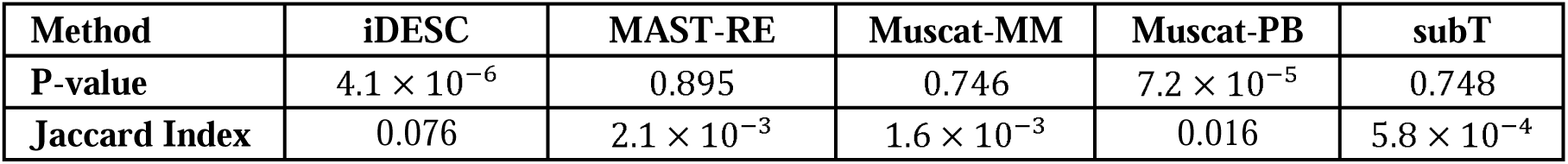
Fisher’s exact test and Jaccard index measuring the DE genes overlapping between two scRNA-seq datasets

| Method | iDESC | MAST-RE | Muscat-MM | Muscat-PB | subT |
| --- | --- | --- | --- | --- | --- |
| **P-value** | $4.1 \times 10^{-6}$ | 0.895 | 0.746 | $7.2 \times 10^{-5}$ | 0.748 |
| **Jaccard Index** | 0.076 | $2.1 \times 10^{-3}$ | $1.6 \times 10^{-3}$ | 0.016 | $5.8 \times 10^{-4}$ |

**Figure 4.**
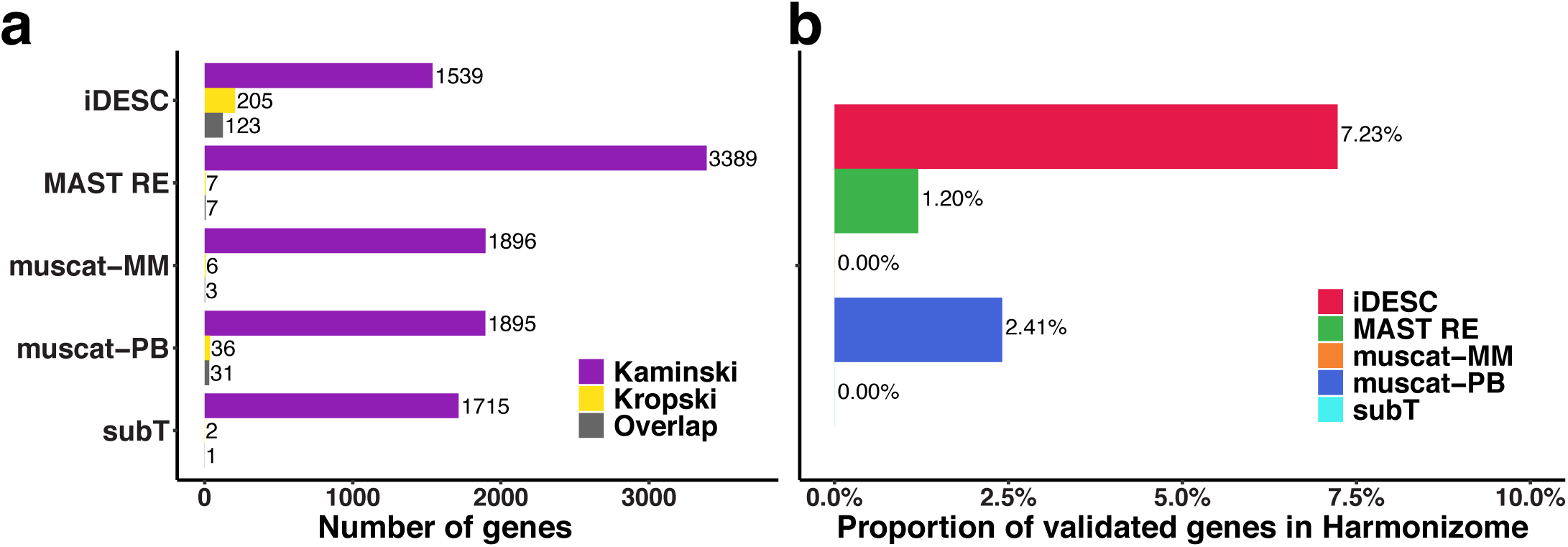
Consistency and validation of DE genes overlapping between two real datasets. A) Barplots show the number of DE genes identified in the Kaminski (purple), Kropski (yellow) datasets and the overlap (grey) between them. B) Barplots show the percentage of IPF-related genes in the Harmonizome database identified in both datasets.

## Discussion

We have developed a new method, iDESC, to detect cell type specific DE genes between two groups of subjects in scRNA-seq data. iDESC fits a zero-inflated GLMM assuming dropouts to have zero count and captured expression to have a negative binomial distribution. Subject effect is modeled as a random effect in the log-mean of the negative binomial component. Wald test is used to assess the group mean difference in captured transcripts. We compared the performance of iDESC with other exiting DE analysis methods. Permutation results demonstrated that the type I error of the methods that consider subject effect were well calibrated, whereas the type I error of the methods that ignore subject effect were highly inflated. Through simulations, we showed that iDESC achieved comparable or higher power to identify DE genes among methods that consider subject effect. In the two IPF scRNA-seq datasets, several DE genes identified by iDESC had been well supported by the literature to be related to IPF pathogenesis. Moreover, iDESC generated the most consistent and validated results between the two datasets. These results demonstrated superior performance of iDESC over the other existing methods we compared, suggesting the importance of considering subject effect and dropouts in the DE analysis of scRNA-seq data with multiple subjects.

Despite the advantages of iDESC over the other DE analysis methods shown in this article, iDESC can be improved in several directions. First, iDESC models gene-specific dropout rate assuming that all cells have the same dropout rate on this gene. A previous study **Error! Reference source not found**. showed that dropout rate may depend on the true underlying gene expression in a cell. Therefore, information across cells can be borrowed and integrated to improve the model of dropout rate. Second, in some cell types, the cell-to-cell heterogeneity of certain genes is so high that negative binomial distribution may not fit the data well. Especially when many cell subtypes are present in the data, the distribution of expression may have multiple modes. Data transformation, a goodness-of-fit test for iDESC and/or replacing negative binomial distribution with a multi-modal distribution or a mixture model may improve the fitting of data. Finally, estimation of dispersion parameter in a negative binomial distribution has been shown to be challenging. Multiple dispersion correction approaches [39, 51-53] that have been developed to improve accuracy can be used to further improve the performance of iDESC.

## Conclusions

We developed iDESC, a zero-inflated negative binomial mixed model that considers both subject effect and dropouts, to identify cell type specific differentially expressed genes in scRNA-seq data with multiple subjects. iDESC had well-calibrated type I error and achieved high sensitivity and specificity in differential expression analysis.

## Materials and methods

### Statistical model

To identify cell type specific DE genes between two groups of subjects, iDESC uses a zero-inflated negative binomial mixed model to consider both subject effect and dropout events in scRNA-seq data with multiple subjects. The model includes two components: a zero component representing dropouts and a negative binomial component representing captured expression. We allow both dropout rate and mean expression to be different between the two groups in the model. Suppose cells are collected from *n* subjects. In a given cell type of interest, subject *i* has *n*_*i*_ cells so that there are in total 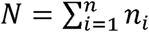 cells of the given type. Let *X*_*i*_ be the group label of subject *i*,where *X*_*i*_ is 1 if subject *i* belongs to group 1 and 0 if subject *i* belongs to group 2. For a given gene, let *Y*_*ij*_ denote the UMI count of this gene in cell *j* from subject *i*. We model the UMI count as:

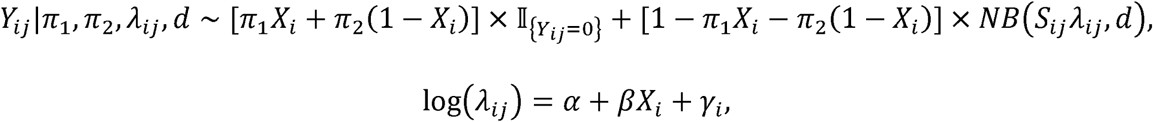

where *π*_1_ and *π*_2_ are the dropout rates in the two groups representing the probability of this gene being dropped out, 𝕀_{°}_ is the indicator function that takes value 1 when the condition in the brackets is satisfied, 0 otherwise, *S*_*ij*_ is the total UMI counts of cell *j* from subject *i, λ*_*ij*_ is the rate parameter of the negative binomial distribution representing the true underlying relative expression level of this gene, and *d* is the dispersion parameter. The rate parameter *λ*_*ij*_ is further modeled as a GLMM with log link, where *α* is the intercept, *β* is the group effect representing the log fold change of mean expression between the two groups, and *γ*_*i*_ is the subject random effect. We assume *γ*_*i*_ are independent and *γ*_*i*_ ∼ *N*(0, *σ*^2^). To test if this gene is differentially expressed between the two groups, we constructed a Wald statistic to test *H*_0_: *β* = 0 against *H*_1_: *β* ≠ 0 using an R package ‘glmmTMB’ [54].

### Real datasets

We evaluated the performance of iDESC and other methods using two scRNA-seq datasets of lung samples from two IPF studies generated using 10X Genomics Chromium platform. Both datasets included IPF patients and healthy controls. In this study, we focused on macrophage which has been recognized to play a significant role in IPF pathogenesis [43-48].

**Kaminski** refers to the scRNA-seq dataset of distal lung parenchyma samples from 32 IPF and 28 control donor lungs in Adams et al. [18]. The raw data include 38,070 genes and 101,230 macrophages.

**Kropski** refers to the scRNA-seq dataset of whole lungs from 12 IPF patients and 10 healthy donors in Habermann et al. [42]. The raw data include 37,647 genes and 11,532 macrophages.

### Data preprocessing

To ensure the same type of macrophages were compared in both datasets so that the DE analysis results were comparable between the Kaminski and Kropski datasets, we compared cell type annotations by integrating these two datasets using the integration analysis in Seurat [55]. Macrophages with substantial overlap in the UMAP of integrated data between the two datasets were extracted for downstream DE analysis. Due to the computational burden in method performance evaluation, we performed down-sampling in both datasets. We first applied a graph-based Louvain clustering algorithm [56] to the integrated data and identified 21 cell clusters in macrophages. Within each cluster, we randomly selected 50 macrophages from each subject if the number of cells for the subject was more than 50 or selected all macrophages on those subjects whose cell number was no more than 50. We then filtered out genes that were expressed in less than 5% of cells on at least one subject from both IPF and control groups. After gene filtering, we ended up with 6,564 genes and 21,433 macrophages from the Kaminski dataset, and 6,128 genes and 10,520 macrophages from the Kropski dataset.

### Type I error assessment

To assess type I error, we randomly permuted the group labels of subjects in both the Kaminski and Kropski datasets so that no model assumptions were made in data generation and the within-subject cell-to-cell correlation structure was preserved in the data. The permuted datasets were not expected to show transcriptomic difference between the two groups. We performed 500 permutations on each of the two datasets and applied all DE analysis methods to the permuted datasets. Genes with p-value < 0.05 were considered to be significant. The empirical type I error for each gene was calculated as the proportion of permuted datasets in which p-value of the gene was less than 0.05.

### Power comparison

To compare the power of the methods, we simulated single-cell expression data from zero-inflated negative binomial mixed models. Library size (total number of UMI counts) and model parameters were set at values estimated from the Kaminski dataset. We considered three parameter settings for the intercept and variance of subject effect, respectively, where they were set at the 25th, 50th and 75th percentile of the estimated values across all genes with *α*= −9.3, −8.9, −8.3 and *σ*^2^ = 0.03, 0.05, 0.08. The dispersion parameter was set to *d* = 0.5, the median of the estimated values across all genes. In the Kaminski dataset, the estimated dropout rates in the disease group were very close to zero and smaller than that in the control group for most genes. So, we designed two scenarios for dropout rates. In the first scenario, dropout rates were the same in the two groups and we set at two levels, *π*_1_ = *π*_2_ =0 which represents no dropout, and *π*_1_ = *π*_2_ = 0.01 which represents a low dropout rate in both groups. In the second scenario, dropout rates were different between the two groups and we set the dropout rate in the disease group as *π*_1_ =0 and that in the control group at two levels with *π*_2_ = 0.01, 0.04, which corresponded to the median and mean of the estimated dropout rate difference between the two groups across all genes. At each simulation setting, we generated single-cell gene expression data containing 500 genes on 60 subjects with 30 in each group and 100 cells per subject. Among the 500 genes, we assumed that 30% were DE genes and the remaining were non-DE genes. For non-DE genes, the group effect was set to *β* = 0, while for the DE genes, *β* was set to values from -0.3 to -0.1, and 0.1 to 0.2 with an increment of 0.05. We applied all methods to the simulated datasets to identify DE genes. The prediction accuracy was measured by the area under an ROC curve (AUC) by comparing the DE genes identified by each method at different p-value threshold with the ground truth.

### scRNA-seq data analysis

We applied all DE analysis methods that consider subject effect to the two IPF scRNA-seq datasets and evaluated their performance based on the consistency of the DE genes between the two datasets. Genes with FDR < 0.1 were considered to be differentially expressed. Cell-level effect coefficient [20] was used to demonstrate subject variation. For each cell, we calculated dot products of log-normalized expression and the estimated group effects across the DE genes identified by iDESC and then scaled to a maximum absolute value of 1. To compare the consistency of the DE genes between the two datasets, Fisher’s exact test was used to assess the significance of the overlap and Jaccard index was calculated to measure the similarity between the two DE gene lists. Jaccard index was calculated as the proportion of overlapping genes among all DE genes identified in either of the datasets. For the overlapping DE genes between the two datasets identified by each method, we validated them using IPF associated genes. We first obtained a list of 83 genes that are related to IPF in the Harmonizome database (https://maayanlab.cloud/Harmonizome/). The proportion of validation was estimated by the percentage of the 83 genes that were identified in both datasets.

## Abbreviations

scRNA-seq: Single-cell RNA sequencing
UMI: Unique Molecular Identifier
IPF: Idiopathic Pulmonary Fibrosis
DMSO: Dimethyl Sulfoxide
ROC: Receiver Operating Characteristic
AUC: Area Under the Curve

## Declarations

### Ethics approval and consent to participate

No ethical approval was required for this study. All public datasets used in the paper were generated by other organizations that have obtained ethical approval.

### Consent for publication

Not applicable

### Availability of data and materials

All analyses were run in R v3.5.3. An R package implementing the proposed method is available at https://github.com/yl883/iDESC. The description of used data sets is in the “Real datasets” section.

### Competing interests

The authors declare that they have no competing interests.

### Funding

This work was supported by National Institutes of Health grants K01AA023321 and R21LM012884, and National Science Foundation grant DMS1916246.

### Authors’ contributions

YL, ZW and XY conceived the idea, developed the method, and designed the study. YL implemented the software and performed the analyses. NW contributed to simulation data collection. TSA, JCS and NK provided IPF lung scRNA-seq data. TSA, JCS, WW, JEM, GLC and NK aided result interpretation. YL, ZW and XY wrote the manuscript. ZW and XY supervised the research. All authors read and approved the final manuscript.

## Acknowledgements

The authors thank Ningshan Li, Qile Dai, Ran Tu and the Kaminski Lab for insightful comments and suggestions that helped improve the quality of the manuscript.

## Notes

https://github.com/yl883/iDESC

### Competing Interest Statement

The authors have declared no competing interest.

## References

1. Gawad, C., W. Koh, and S.R. Quake, Single-cell genome sequencing: current state of the science. Nat Rev Genet, 2016. 17(3): p. 175–88.

2. Macosko, E.Z., et al., Highly Parallel Genome-wide Expression Profiling of Individual Cells Using Nanoliter Droplets. Cell, 2015. 161(5): p. 1202–1214.

3. Zheng, G.X., et al., Massively parallel digital transcriptional profiling of single cells. Nat Commun, 2017. 8: p. 14049.

4. Stephenson, W., et al., Single-cell RNA-seq of rheumatoid arthritis synovial tissue using low-cost microfluidic instrumentation. Nature Communications, 2018. 9.

5. Segerstolpe, A., et al., Single-Cell Transcriptome Profiling of Human Pancreatic Islets in Health and Type 2 Diabetes. Cell Metab, 2016. 24(4): p. 593–607.

6. Yao, C., et al., Single-cell RNA-seq reveals TOX as a key regulator of CD8(+) T cell persistence in chronic infection. Nature Immunology, 2019. 20(7): p. 890-+.

7. Pop, S.M., et al., Single cell analysis shows decreasing FoxP3 and TGF beta 1 coexpressing CD4(+)CD25(+) regulatory T cells during autoimmune diabetes. Journal of Experimental Medicine, 2005. 201(8): p. 1333–1346.

8. Chung, W., et al., Single-cell RNA-seq enables comprehensive tumour and immune cell profiling in primary breast cancer. Nat Commun, 2017. 8: p. 15081.

9. Vieira Braga, F.A., et al., A cellular census of human lungs identifies novel cell states in health and in asthma. Nat Med, 2019. 25(7): p. 1153–1163.

10. Grun, D., et al., Single-cell messenger RNA sequencing reveals rare intestinal cell types. Nature, 2015. 525(7568): p. 251-+.

11. Buettner, F., et al., Computational analysis of cell-to-cell heterogeneity in single-cell RNA-sequencing data reveals hidden subpopulations of cells. Nat Biotechnol, 2015. 33(2): p. 155–60.

12. Yuan, G.C., et al., Challenges and emerging directions in single-cell analysis. Genome Biol, 2017. 18(1): p. 84.

13. Shalek, A.K. and M. Benson, Single-cell analyses to tailor treatments. Sci Transl Med, 2017. 9(408).

14. Luecken, M.D. and F.J. Theis, Current best practices in single-cell RNA-seq analysis: a tutorial. Mol Syst Biol, 2019. 15(6): p. e8746.

15. Squair, J.W., et al., Confronting false discoveries in single-cell differential expression. Nature Communications, 2021. 12(1): p. 5692.

16. Kharchenko, P.V., L. Silberstein, and D.T. Scadden, Bayesian approach to single-cell differential expression analysis. Nat Methods, 2014. 11(7): p. 740–2.

17. Reyfman, P.A., et al., Single-Cell Transcriptomic Analysis of Human Lung Provides Insights into the Pathobiology of Pulmonary Fibrosis. Am J Respir Crit Care Med, 2019. 199(12): p. 1517–1536.

18. Adams, T.S., et al., Single-cell RNA-seq reveals ectopic and aberrant lung-resident cell populations in idiopathic pulmonary fibrosis. Sci Adv, 2020. 6(28): p. eaba1983.

19. Zimmerman, K.D., M.A. Espeland, and C.D. Langefeld, A practical solution to pseudoreplication bias in single-cell studies. Nature Communications, 2021. 12(1).

20. Crowell, H.L., et al., muscat detects subpopulation-specific state transitions from multi-sample multi-condition single-cell transcriptomics data. Nat Commun, 2020. 11(1): p. 6077.

21. Lopez, R., et al., Deep generative modeling for single-cell transcriptomics. Nat Methods, 2018. 15(12): p. 1053–1058.

22. Wohnhaas, C.T., et al., DMSO cryopreservation is the method of choice to preserve cells for droplet-based single-cell RNA sequencing. Sci Rep, 2019. 9(1): p. 10699.

23. Soneson, C. and M.D. Robinson, Bias, robustness and scalability in single-cell differential expression analysis. Nat Methods, 2018. 15(4): p. 255–261.

24. Dal Molin, A., G. Baruzzo, and B. Di Camillo, Single-Cell RNA-Sequencing: Assessment of Differential Expression Analysis Methods. Front Genet, 2017. 8: p. 62.

25. Jaakkola, M.K., et al., Comparison of methods to detect differentially expressed genes between single-cell populations. Briefings in Bioinformatics, 2017. 18(5): p. 735–743.

26. Vallejos, C.A., J.C. Marioni, and S. Richardson, BASiCS: Bayesian Analysis of Single-Cell Sequencing Data. PLoS Comput Biol, 2015. 11(6): p. e1004333.

27. Jia, C., et al., Accounting for technical noise in differential expression analysis of single-cell RNA sequencing data. Nucleic Acids Res, 2017. 45(19): p. 10978–10988.

28. Qiu, X., et al., Single-cell mRNA quantification and differential analysis with Census. Nat Methods, 2017. 14(3): p. 309–315.

29. Qiu, X., et al., Reversed graph embedding resolves complex single-cell trajectories. Nat Methods, 2017. 14(10): p. 979–982.

30. Trapnell, C., et al., The dynamics and regulators of cell fate decisions are revealed by pseudotemporal ordering of single cells. Nat Biotechnol, 2014. 32(4): p. 381–386.

31. Chen, W., et al., UMI-count modeling and differential expression analysis for single-cell RNA sequencing. Genome Biol, 2018. 19(1): p. 70.

32. Miao, Z., et al., DEsingle for detecting three types of differential expression in single-cell RNA-seq data. Bioinformatics, 2018. 34(18): p. 3223–3224.

33. Ye, C., T.P. Speed, and A. Salim, DECENT: differential expression with capture efficiency adjustmeNT for single-cell RNA-seq data. Bioinformatics, 2019. 35(24): p. 5155–5162.

34. Wu, Z., et al., Two-phase differential expression analysis for single cell RNA-seq. Bioinformatics, 2018. 34(19): p. 3340–3348.

35. Finak, G., et al., MAST: a flexible statistical framework for assessing transcriptional changes and characterizing heterogeneity in single-cell RNA sequencing data. Genome Biol, 2015. 16: p. 278.

36. Vieth, B., et al., A systematic evaluation of single cell RNA-seq analysis pipelines. Nat Commun, 2019. 10(1): p. 4667.

37. Korthauer, K.D., et al., A statistical approach for identifying differential distributions in single-cell RNA-seq experiments. Genome Biol, 2016. 17(1): p. 222.

38. Delmans, M. and M. Hemberg, Discrete distributional differential expression (D3E)--a tool for gene expression analysis of single-cell RNA-seq data. BMC Bioinformatics, 2016. 17: p. 110.

39. Love, M.I., W. Huber, and S. Anders, Moderated estimation of fold change and dispersion for RNA-seq data with DESeq2. Genome Biol, 2014. 15(12): p. 550.

40. Ritchie, M.E., et al., limma powers differential expression analyses for RNA-sequencing and microarray studies. Nucleic Acids Res, 2015. 43(7): p. e47.

41. Wilcoxon, F., Individual comparisons of grouped data by ranking methods. J Econ Entomol, 1946. 39: p. 269.

42. Habermann, A.C., et al., Single-cell RNA sequencing reveals profibrotic roles of distinct epithelial and mesenchymal lineages in pulmonary fibrosis. Sci Adv, 2020. 6(28): p. eaba1972.

43. Wynes, M.W. and D.W. Riches, Transcription of macrophage IGF-I exon 1 is positively regulated by the 5’-untranslated region and negatively regulated by the 5’-flanking region. Am J Physiol Lung Cell Mol Physiol, 2005. 288(6): p. L1089–98.

44. Morse, C., et al., Proliferating SPP1/MERTK-expressing macrophages in idiopathic pulmonary fibrosis. European Respiratory Journal, 2019. 54(2).

45. Wang, H., et al., Bioinformatics analysis on differentially expressed genes of alveolar macrophage in IPF. Experimental lung research, 2019. 45(9-10): p. 288–296.

46. Bargagli, E., et al., Macrophage-derived biomarkers of idiopathic pulmonary fibrosis. Pulmonary medicine, 2011. 2011.

47. Schupp, J.C., et al., Macrophage activation in acute exacerbation of idiopathic pulmonary fibrosis. PloS one, 2015. 10(1): p. e0116775.

48. Prasse, A., et al., CCL18 as an indicator of pulmonary fibrotic activity in idiopathic interstitial pneumonias and systemic sclerosis. Arthritis & Rheumatism, 2007. 56(5): p. 1685–1693.

49. Wan, H., et al., Identification of Hub Genes and Pathways Associated With Idiopathic Pulmonary Fibrosis via Bioinformatics Analysis. Frontiers in molecular biosciences, 2021: p. 790.

50. Rouillard, A.D., et al., The harmonizome: a collection of processed datasets gathered to serve and mine knowledge about genes and proteins. Database-the Journal of Biological Databases and Curation, 2016.

51. Robinson, M.D. and G.K. Smyth, Small-sample estimation of negative binomial dispersion, with applications to SAGE data. Biostatistics, 2008. 9(2): p. 321–332.

52. Lloyd-Smith, J.O., Maximum Likelihood Estimation of the Negative Binomial Dispersion Parameter for Highly Overdispersed Data, with Applications to Infectious Diseases. Plos One, 2007. 2(2).

53. Rao, C.R., Large Sample Tests of Statistical Hypotheses Concerning Several Parameters with Applications to Problems of Estimation. Proceedings of the Cambridge Philosophical Society, 1948. 44(1): p. 50–57.

54. Brooks, M.E., et al., glmmTMB Balances Speed and Flexibility Among Packages for Zero-inflated Generalized Linear Mixed Modeling. R Journal, 2017. 9(2): p. 378–400.

55. Stuart, T., et al., Comprehensive Integration of Single-Cell Data. Cell, 2019. 177(7): p. 1888–1902 e21.

56. Blondel, V.D., et al., Fast unfolding of communities in large networks. Journal of Statistical Mechanics-Theory and Experiment, 2008.

