## Supplementary figures and images for "iDESC: Identifying differential expression in single-cell RNA sequencing data with multiple subjects"

### Supplementary Figure 1

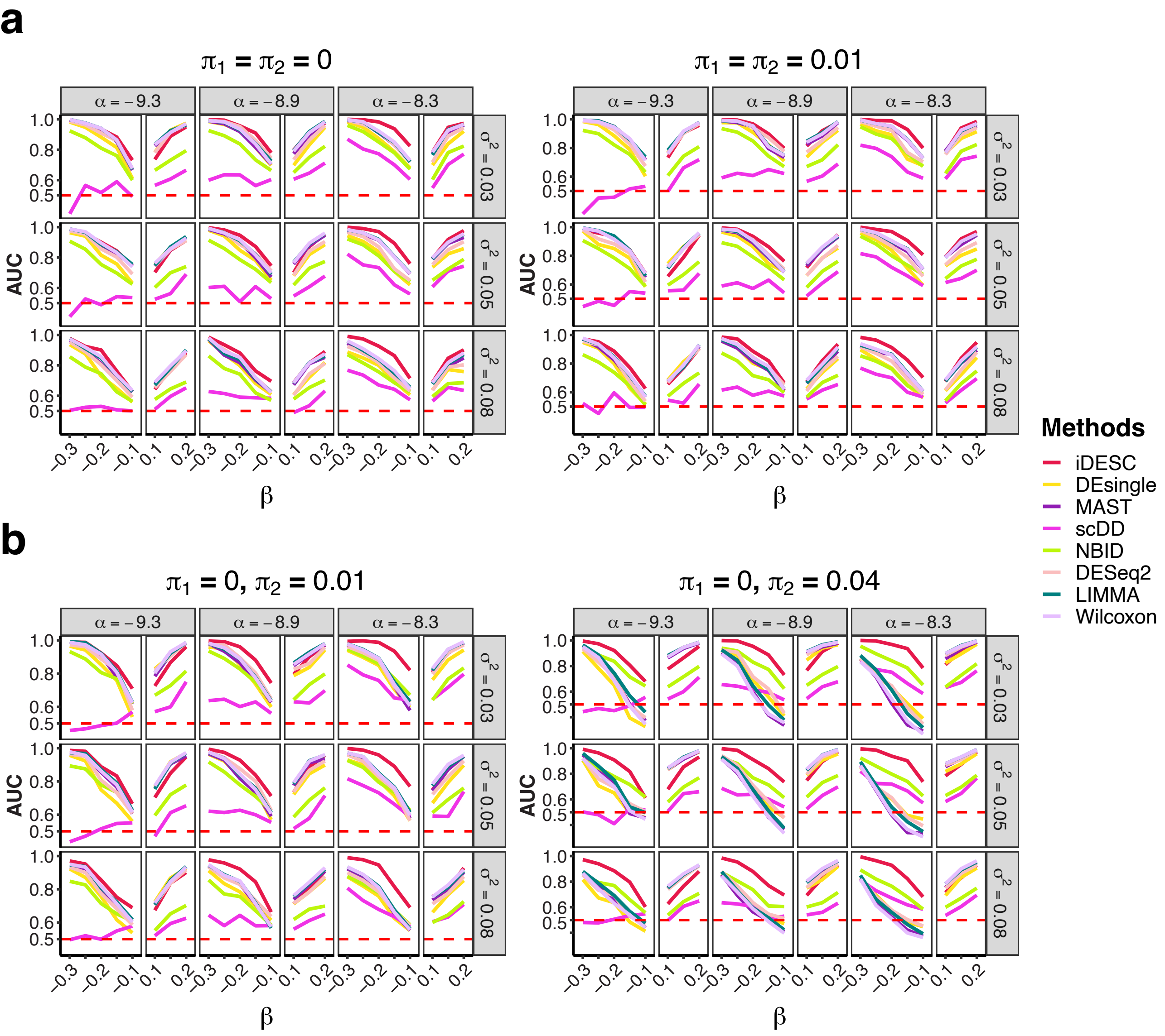
